# Intermittent hormonal therapy shows similar outcome than SOC in ER+ breast cancer preclinical model

**DOI:** 10.1101/509158

**Authors:** Pedro M. Enriquez-Navas, Libia Garcia, Mahmoud Abdalah, Olya Stringfield, Kimberly Luddy, Sabrina Hassan, Robert J. Gillies, Robert A. Gatenby.

**Author notes:** Corresponding Author Pedro M. Enriquez-Navas.

## Abstract

Clinical breast cancers in which at least 10% of cells express the estrogen receptor are labeled as “ER positive.” First line therapy for these patients is typically continuous administration of anti-estrogen drugs at maximum tolerated dose (MTD) until progression. In the vast majority of patients, resistance to hormone therapy evolves in the breast cancer cells within 2 years leading to treatment failure and tumor progression. In prior studies, we have demonstrated continuous application of MTD chemotherapy results in evolutionary dynamics (termed “competitive release”) that accelerates proliferation of treatment-resistance populations. In contrast, evolution-informed application of treatment reduces drug administration to maintain substantial populations of therapy-sensitive cells to reduce proliferation of resistant phenotypes. Prior pre-clinical and clinical studies have shown this strategy can delay or prevent proliferation of resistant cells and prolong time to progression (TTP). We hypothesize that similar dynamics may be observed in hormonal therapy of ER+ breast cancers. Here we address two important dynamics. First, we consider a clinical scenario in which symptoms are sufficiently severe or life-threatening to require rapid and substantial tumor reduction. Can this be achieved while retaining evolutionary dynamics to subsequently delay proliferation of resistance? A second, related question is defining the cost of resistance to anti-estrogen therapy. Here, we investigated the evolutionary dynamics of resistance to anti-estrogen therapy using ER+ MCF-7 orthotropic xenografts treated with both continuous Tamoxifen as well as cycles in which estrogen stimulation is combined with estrogen suppression. As expected, continuous administration of anti-estrogen drugs successfully suppressed tumor growth. However we found that brief interruptions in drug administration permitted equal tumor control while administering up to 50% less drug and maintaining cell phenotypes that retained high levels of ER expression and lower levels of MDR1 expression. In follow-on experiments combining hormonal and chemo-therapies; we obtained similar tumor control to hormonal therapy alone but with more necrosis and significantly lower ER expression in the surviving population.

## Introduction

In 2017 about 300,000 women in the United States were diagnosed with breast cancer and, among these, almost 90% were characterized as estrogen receptor positive (ER+) [1, 2]. The criteria for ER+ requires that at least 10% of the tumor cells express ER on immunohistochemical staining so that, in many breast cancers, a significant fraction of the cancer cells do not express ER and may be resistant to anti-estrogen therapy [3].

Initial treatment for ER+ breast cancer includes blockade of the effect of estrogen by selective estrogen receptor modulator (SERM) drugs.[4] Tamoxifen, which blocks the interaction between estrogen and its receptor impeding cell replication, is among the most widely used SERM drugs and is typically administered at maximum tolerated dose daily until tumor progression [5, 6]. Alternative strategies, such as aromatase inhibitors, block the synthesis of estrogen by normal cells. While nearly all ER+ tumors initially respond to anti-estrogen therapy, evolution of resistance with treatment failure and tumor progression typically is observed within a few months to a few years[7].

There are three major drawbacks in anti-estrogen treatment: (1) cost and side effects reduce compliance in up to 25% of the patients [8]; (2) about 10% will develop one or more side-effects that require dose adjustments or treatment cessation [1, 7, 9]; (3) prolonged continuous treatment may significantly increase risk for endometrial cancer. Nevertheless, the vast majority of patients initially respond to anti-estrogen therapy but development of resistance leading to treatment failure and progression is virtually inevitable [10–12] and, among the multiple mechanisms of resistance, evolution of estrogen-independent growth is most common[11].

A common evolution-based strategy to delay tumor progression focuses on the phenotypic cost of resistance. This is readily apparent in resistance to chemotherapy based on increased expression of MDR1 (Multi-Drug Resistance system-1); a membrane glycoprotein (PgP) that is an active ATPase pump extruding lipophilic cationic xenobiotics. In some studies, up to 40% of a cell’s energy budget must be used to synthesize, maintain, and operate MDR membrane pumps – an obvious cost of resistance [13, 14]. Thus, a strategy termed “adaptive therapy” explicitly limits cancer treatment to maintain a significant population of treatment-sensitive cells. Therapy is then withdrawn. However, in the subsequent tumor regrowth, the sensitive cells, in the absence of the cost of resistance, outcompete the resistant cells. Thus, through multiple cycles, the tumor population remains sensitive to the primary treatment. In an ongoing clinical trial in metastatic castrate-resistant prostate cancer, we have found that treatment that only reduces the serum PSA to half of its pre-treatment value can both substantially increase the time to progression while decreasing the cumulative drug does [15].

A number of questions regarding optimal evolution-based treatments remain. Among these, perhaps the most urgent is the apparently conflicting demands for treatment in which a patient presents with highly symptomatic or potentially life-threatening conditions. Here, rapid and significant reduction of the tumor burden is clinically necessary but could also result in competitive release of resistant clones that result in rapid proliferation leading to tumor failure and tumor progression with recurrent symptoms. Here we examine potential treatment strategies that can both rapidly diminish tumor burden to very low levels while maintaining evolutionary dynamics that can prolong tumor control and reduce the cumulative drug dose to reduce toxicity and cost.

A second question in this study is the cost of resistance to hormonal therapy. Although drug efflux by the MDR proteins is a mechanism of resistance in ER+ cells, clinical studies have found that, in general, durable resistance to estrogen therapy is most commonly obtained in breast cancer cells through expression of alternative pathways that permit estrogen-independent survival and proliferation. However, unlike the dynamics of PgP, the evolutionary cost of estrogen-independence is not obvious. Nevertheless, that such a cost exists can be inferred using a concept termed “evolutionary triage.” Briefly, “evolutionary triage”[15] simply states that, among competing populations, the fittest phenotype will be the most proliferative and, in general, be the largest population. Therefore, in general, the relative fitness of each cancer subpopulation can be estimated by their relative abundance within the tumor. This, however, yields puzzling results for ER+ breast cancers in which greater than 50% of the cells do not express the ER on immunohistochemical stains. Despite ER+ cells being in the minority, these tumors still typically respond, at least initially, to anti-estrogen therapy suggesting some component of the treatment dynamics is not being captured in the IHC results. Thus, it is not clear if evolution-based treatment strategies could be successfully applied in clinical treatment of ER+ metastatic breast cancer.

Here, we address these questions in pre-clinical studies. Our results showed that (i) intermittent therapies can control tumor growth with less Tamoxifen (using as little as 50% of the standard dose), (ii) the ER expression is maintained at or above the levels of SOC, (iii) expression of MDR1 was reduced in tumors treated with intermittent tamoxifen therapy, and (iv) the combination of hormonal- and chemo-n the standard therapy, although showing more tumor necrosis.

We conclude that evolution-based administration of anti-estrogen drugs in patients is likely to benefit patients with metastatic ER+ breast cancer compared to current strategies of continuous MTD dosing until progression. Our results, however, also suggest the evolutionary dynamics that govern estrogen-related fitness in breast cancer cells and the clinical efficacy of anti-estrogen therapy are not fully understood and require further investigation.

## Materials and Methods

### In-vivo Experiments

Different cohorts (n= 10, 14, 4, and 39) of nude (nu/nu) mice were injected in the mammary fat pad with 5*10^6^ MCF7 cells (injected in a mixture 1:1 with phenol-red free Matrigel) tagged with GFP (cells were obtained from ATCC, and were grown following its guidelines, cell culture media and supplies were obtained from Thermo Fisher Scientific). Prior to the cell injections, an estrogen pellet (Innovative Research of America) of 0.72 mg of β-estrogen with 90 days slow release was subcutaneously implanted in each mouse, giving a continuous dose of 400 pg of estrogen per milliliter of blood. If the experiments lasted longer than 90 days, a similar pellet was implanted to the mice.

When tumors reached 300 mm^3^ approximately, mice were randomly distributed in groups and treated following one of the subsequent treatments: *Controls* (Ctrl), no treatment; *Tamoxifen standard* (TamST), 0.5 mg of tamoxifen per mouse daily; *Tamoxifen-vacation* (TamVac), 2 weeks of TamST followed by one week of no treatment (vacation); *Tamoxifen 2 weeks* (Tam2weeks), two weeks of TamST followed by 2 weeks of vacation; *Tamoxifen 3 weeks* (Tam3weeks), three weeks of TamST followed by three weeks of vacation; *Tamoxifen and Paclitaxel* (TamPac), 2 weeks of TamST followed by one week in which Paclitaxel is applied in two non-consecutive days at a concentration of 20mg/kg (ip) (PacST); *Vacation and Paclitaxel* (VacPac), 2 weeks of no treatment followed by one week in which 20 mg/kg of Paclitaxel in applied in two non-consecutive days; *Paclitaxel and Tamoxifen* (PacTam), one week of PacST followed by two weeks of TamST.

Tamoxifen (Caiman Chemical Company) was suspended in peanut oil (Sigma Aldrich) and given to animals by either i.p or gavage routes; initially Tamoxifen was administered in 200 µl of peanut oil through i.p. injection. However, we changed the route of administration to gavage because mice were not able to metabolize the peanut oil and all of them died in a short period of time; maybe due to peritonitis (when tumors were collected, we noticed that peanut oil was accumulated in mice abdomen). Paclitaxel (LC laboratories) was dissolved in a mixture of Koliphor oil and ethanol (1:1, both solvents were obtained from Sigma Aldrich) and it was given via i.p. injection; before injection, the corresponding dose was diluted twice with PBS.

Mice were visually monitored, and their weights monitored once a week, to address any treatment toxicity issues. Tumor growth was measured once a week by either caliper, using the formula: 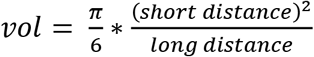, or by MRI; magnetic resonance data were acquired with a 7 T horizontal magnet Agilent ASR 310 (Agilent Technologies Inc.) equipped with nested 205/120/HDS gradient insert and a bore size of 310 mm. Before imaging, the animals were placed in an induction chamber and anesthetized with 2-3% isoflurane delivered in 1.5 liter/min oxygen ventilation. After complete induction, animals were restrained in a custom-designed holder and inserted into the magnet while constantly receiving isoflurane (1 to 3%) within the same oxygen ventilation. Body temperature (37° ± 1°C) and respiratory functions were monitored continuously (SAII System) during the experimental time. A 35 mm Litzcage coil (Doty Scientific) was used to carry out axial T2-weighted fast spin-echo multislice experiments (acquired with TE/TR [echo time/repetition time] = 72 ms/1000 ms, field of view (FOV) = 35 × 35 mm^2^, matrix = 128 × 128, yielding a spatial in-plane resolution of 273 µm, slice thickness of 1.5 mm).

At the completion of the experiments (either tumor under control or when tumor volume reached ~2000 mm^3^) tumors were collected for histological analysis. After collection, tumors were processed for histological studies by soaking in formalin, during 24 h at least, followed by embedding in paraffin blocks. Consecutive histological slices (5 µm thickness) were cut from each tumor to study the necrosis percentage (H&E), vascularity density and functionality (CD31 and SMA, respectively), estrogen receptor (ER) expression, and resistance mechanisms (by means of MDR1 expression). Once stained, the slices were imaged at the Moffitt Cancer Center microscope core facilities, using Aperio ScanScope XT microscope and Aperio Spectrum version 10.2.5.2352 image analytic software (Leica Biosystems Inc.). To optimize the image analysis, we trained the analytic algorithm by using ROIs that were selected manually to represent ROIs that are positive or negative for each stain. After initial algorithm training, the software developed a final algorithm, which was used to automatically analyze the slides with a pixel-size resolution (5 µm × 5 µm).

### Imaging analysis

To analyze the homogeneity in the ER expression, a radiomics analysis was performed on IHC estrogen receptor images. Features extraction was done on IHC digital images with the same magnification (x20). Color images were segmented using a thresholding method to identify the pixels that were positively stained. Pixels outside the segmented range (background and/or unstained cells) were discarded and not used for analyses. Masks from positively stained cells were used to generate neighborhood maps. For each pixel in the mask, the number of immediate neighbors that were also in the mask (positively stained) was counted (up to 8) and that number is the pixel’s neighborhood coefficient. Thus, for all samples, the masks of positively stained pixels were replaced by corresponding neighborhood maps consisting of neighborhood coefficients. The neighborhood maps were computed to capture the distribution and the density of positively (ER+) stained pixels.

Following the generation of neighborhood maps, 202 2D image features were calculated from them, including statistical, shape, and texture variables. These features were then reduced by including only one feature from subsets of inter-correlated features (Pearson correlation coefficient > |0.8|). Among the final features list, only those features correlated with heterogeneity in the tissue texture were used to perform the analysis; table 1 shows a brief description of these features. All the animal work during this project was done following the IACUC regulations of University of South Florida (Tampa) at the Moffitt Cancer Center facilities.

Statistical calculations were performed using the Excel software. Student’s t-tests were performed considering two-tailed distribution and two samples with unequal variance.

## Results

This project was designed to understand the evolutionary dynamics of resistance in ER+ breast tumors and, consequently, to improve first-line treatment of estrogen receptor positive (ER+) breast cancers with SERMs, such as Tamoxifen. Treatment algorithms used in this study were suggested by preliminary *in-vitro* data in which MCF7 cells were grown under different microenvironment conditions to study the expression of the estrogen receptor following addition of Tamoxifen and/or Paclitaxel media.

In the first cohort of mice (n = 10) bearing orthotopic MCF7 tumors, Tamoxifen treatment was administered by i.p. injections. As it is shown in **Figure 1A**, treatment algorithm with a one-week vacation (TamVac) maintained tumors at similar volumes as standard treatment (TamST). No increase in tumor volume was observed during the vacation period. Furthermore, at completion of the study, the remaining tumor cells demonstrated greater ER expression compared to the control and equal or greater expression the continuous dose cohort **(Figure 2A)**.

**Figure 1:**
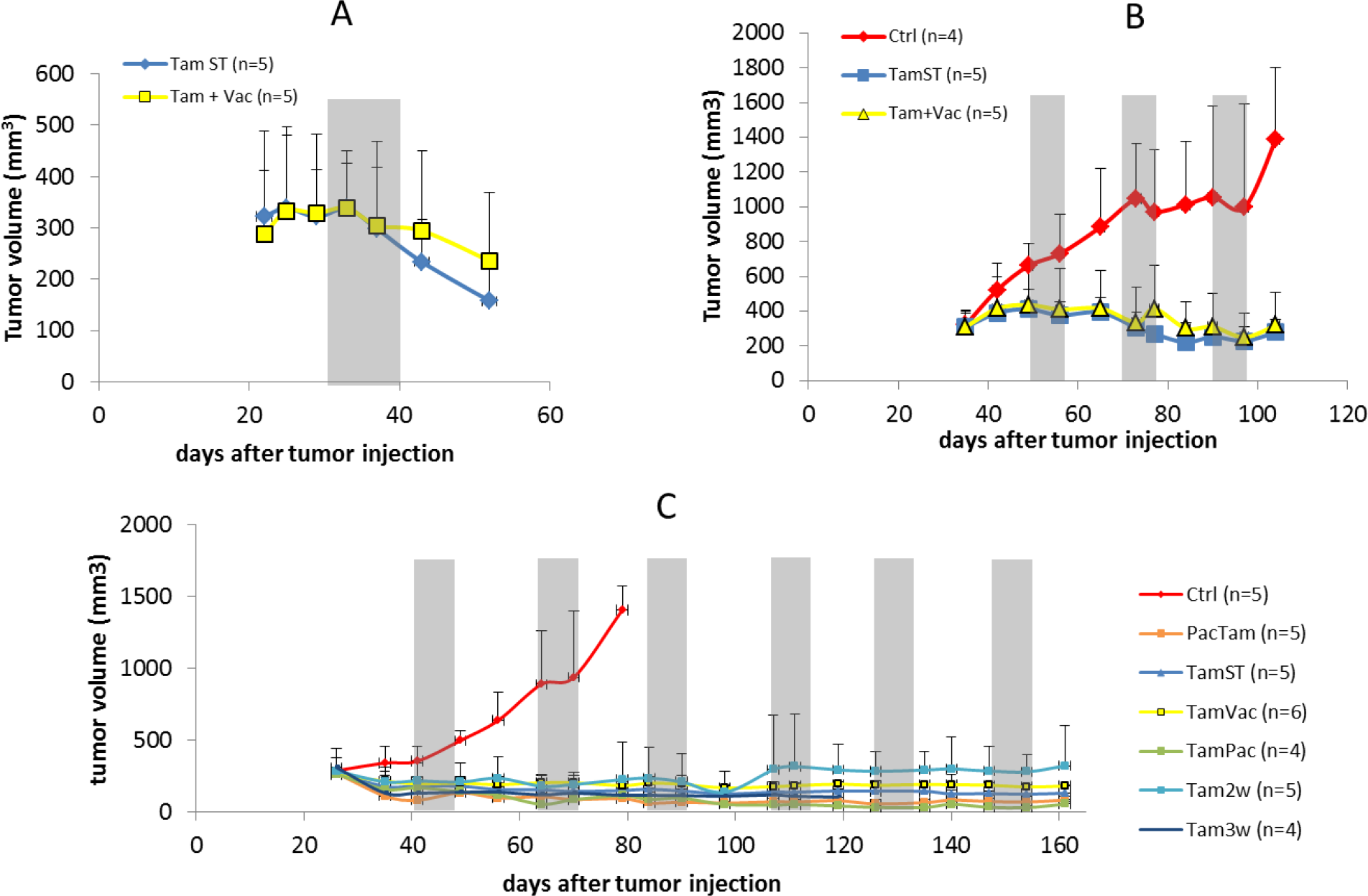
Tumor volumetric data of mice under different treatments. Grey zones correspond to either vacation periods or paclitaxel application (more information about each treatment can be found in Material and Methods section). Tumor volumes were measured by MRI. In parenthesis is the number of mice in each group. Data are shown by mean and error bars represent the standard deviation.

**Figure 2:**
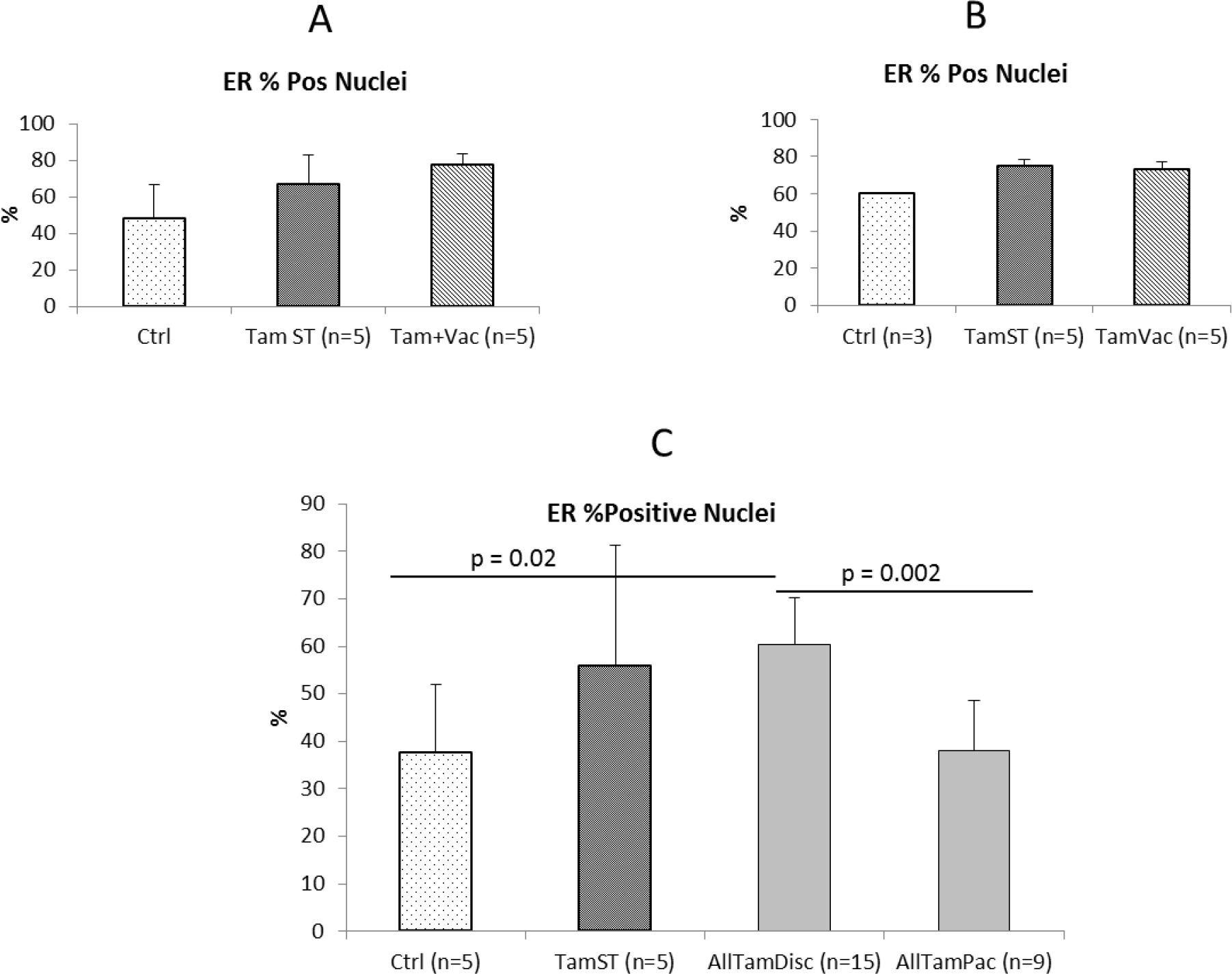
IHC analysis of the ER expression under different treatments. (Significance calculations were done by using a Student t-test with two-tailed distribution and considering two samples with unequal variance).

TamST treatment achieved the same level of control as continuous high dose tamoxifen. This is in contrast to prior reports in which this combination was unsuccessful. [16–18]. It is possible this difference is due to treatment schedule. We treated the animals 7 days per week (matching the daily dose used clinically) and it appears that in the prior studies treatment was not administered on weekends. While the results were encouraging, we noted that all of the mice developed increased peritoneal fluid that appeared to be caused by the peanut oil used in the Tamoxifen injections.

In a second cohort (n=14) we used higher concentration of Tamoxifen (50 mg/ml) with 20 µl of the suspension injected i.p. daily. This reduced the peritoneal fluid collection and showed the same outcome as the prior cohort (**Figure 1B**). Histological analysis of ER expression showed no significant difference between the intermittent and standard tamoxifen therapies (**Figure 2B**). However, the vascularity (density and functionality) was decreased in TamVac therapy (**Figure 3**, columns A and B). MDR1 was expressed in the Tamoxifen therapy groups suggesting membrane extrusion play a significant role in evolution of resistance in this setting (**Figure 4B**).

**Figure 3:**
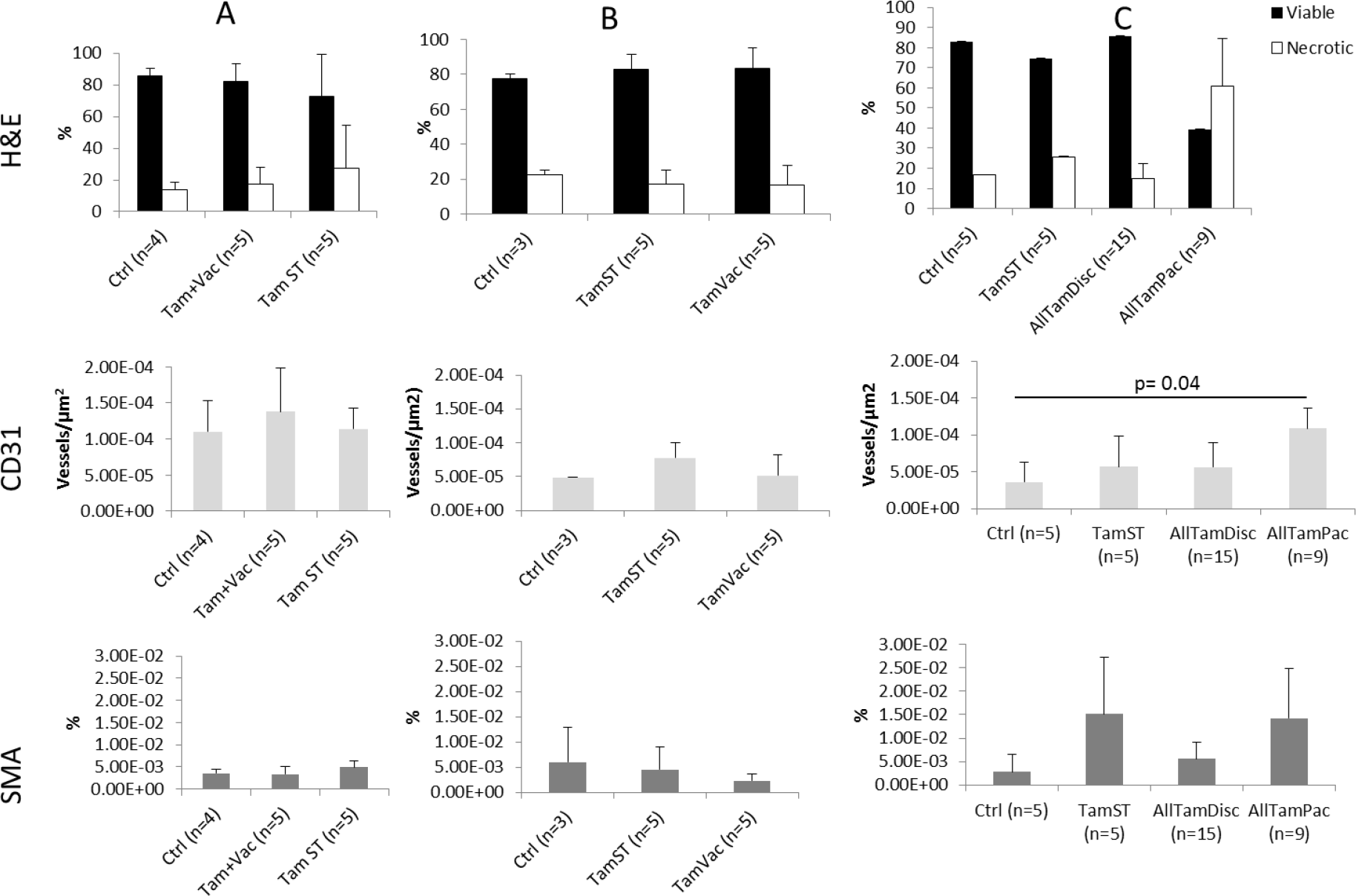
IHC analysis of viable and necrotic tissues (H&E), vessel density (CD31) and functionality (SMA) in tumors under different treatments. Data are shown by mean values with the standard deviation (error bars). p value calculated using Student t-test with two-tailed and unequal variance.

**Figure 4:**
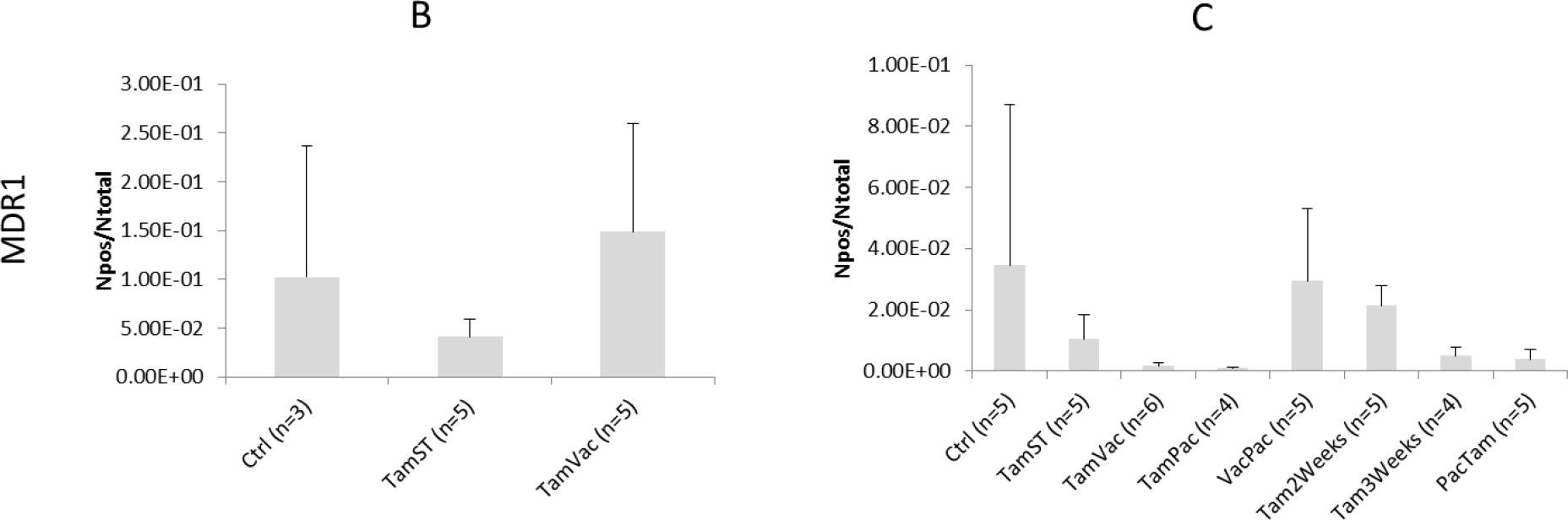
IHC analysis of the expression of MDR1 systems for cohorts B and C. Data is shown by mean with the standard deviation (error bars).

Finally, because the ip injection of Tamoxifen was associated with peritoneal lipid collections, we examined an additional cohort of mice (n=41) in which Tamoxifen was administered by gavage (no significant difference was found between ip and gavage treatments, **Sup. Figure 1**). This cohort included prior treatment (continuous Tamoxifen dosing and one-week vacation period) but also examined longer “vacation” periods during which treatment was suspended for 2 or 3 weeks (Tam2weeks and Tam3weeks). We also examined alternative sequences with Tamoxifen first followed by Paclitaxel or vice-versa (TamPac and PacTam, respectively).

After 130 days of treatment, no significant differences in tumor control were noted in the groups (**Figure 1C**). However, TamVac, Tam2weeks, and Tam3weeks groups had a cumulative dose reduction of 33, 50, and 50%, respectively. No significant tumor growth during these vacation periods was noted (**Figure 1** (grey zones represents vacations periods)). At necropsy, the tumors treated with vacation periods had slightly higher expression of estrogen receptor compared to continuous Tamoxifen (**Figure 2C and Sup. Figure 2**). Interestingly, vessel density and functionality were increased in tumors in which hormonal therapy was combined with a cytotoxic drug (**Figure 3**) but these tumors also demonstrated relatively larger fractions of necrosis (**Figure 3** and **Sup. Figure 3**) when compared to the other cohorts. We studied the expression of MDR1 systems for cohorts B and C using immunohistochemistry (IHC) (**Figure 4**). In general, cohorts with vacation periods showed lower expression of MDR1 than the continuous Tamoxifen group (**Figure 4**, B and C) reflecting the diminished selection pressure for resistance during treatment vacation.

We analyzed the homogeneity in the ER expression in the treatment cohorts using a “neighborhood” imaging analysis, which examined variations in ER expression in physically adjacent cell groups. To do so, we created a mask for the tumor slices following IHC staining for ER expression. An algorithm calculates probability that each ER+ pixels will have similar adjacent pixels (up to 8). This analysis found increased numbers of ER+ similar “neighbors” in treated tumors with either Tamoxifen SOC or intermittent therapies compared to tumors also treated with chemotherapy (**Figure 5**).

**Figure 5:**
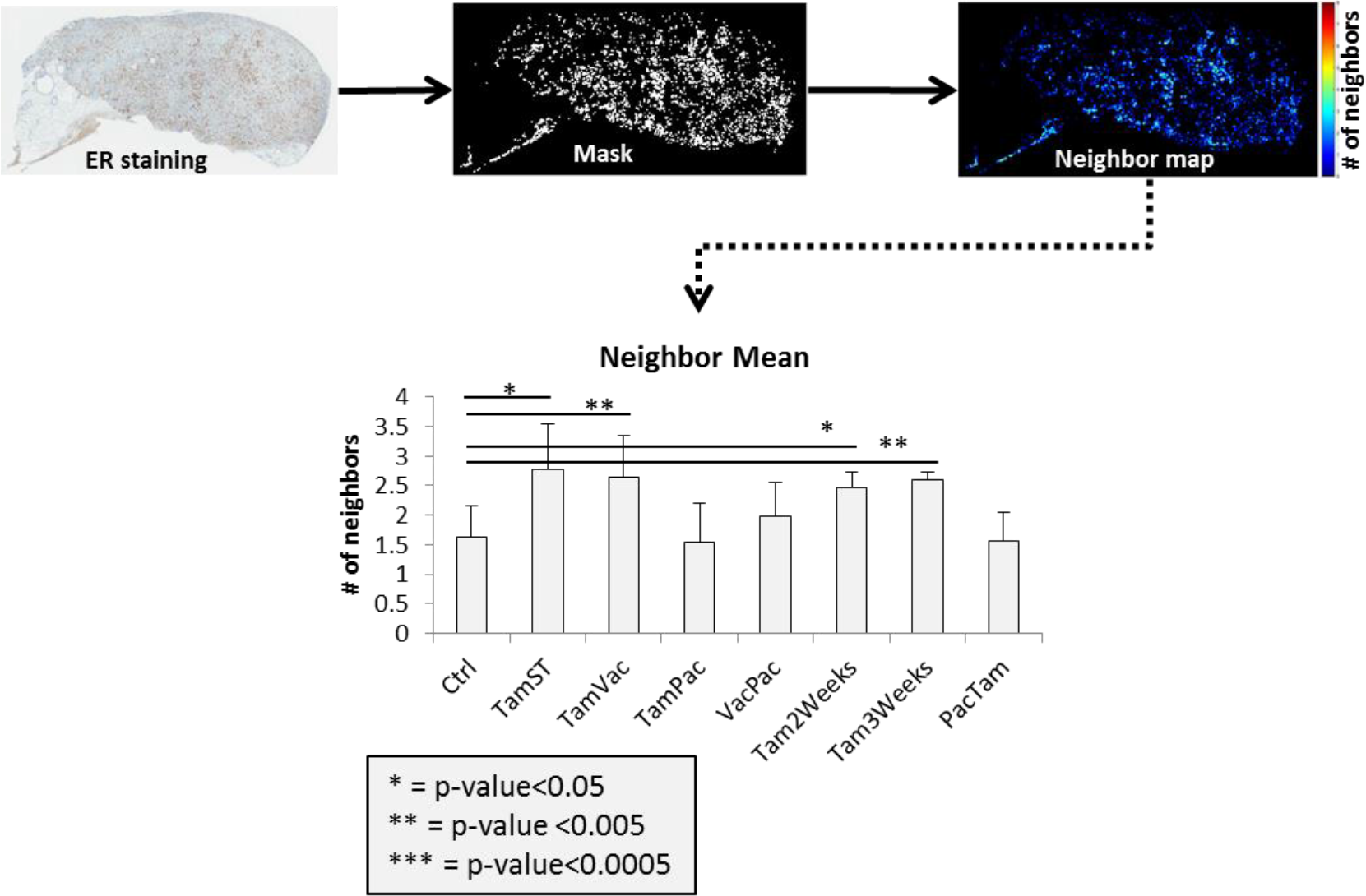
Radiomics analysis of ER expression. The neighbor map (ER+ pixels surrounded by similar ones) showing the mean value. Mean±SD is represented in bar graph. p-values are shown to compare the treated groups with the control (untreated) one.

## Discussion

Cancer cells, like all living systems, evolve to adapt to local environmental selection forces. When clinical therapy is applied, evolution of resistance is commonly observed leading to treatment failure and tumor progression. Here we examine the evolutionary dynamics of ER+ breast cancer treated with anti-estrogen therapy, the typical first line clinical treatment for ER+ breast cancers. In prior pre-clinical and theoretical studies we have found that continuous application of therapy at MTD maximally selects for resistance – a well-known phenomenon in pest management termed “competitive release.” By periodically withdrawing therapy, we reduced the environmental selection forces for resistance by permitting survival of some treatment-sensitive cells. In the absence of treatment, the fitness advantage of the sensitive cells tended to suppress proliferation of the resistant phenotype thus prolonging tumor response. Here we addressed two potential barriers for applying this strategy to ER+ clinical breast cancers. First, in patients who are symptomatic, optimal therapy must reduce the tumor burden below some symptomatic threshold before it is withdrawn. Second, we were concerned with the possibility that the tumor might rapidly progress after treatment withdrawal leading to rapid loss of control and thus decreased TTP.

In these pre-clinical experiments with ER+ breast cancers, we demonstrated intermittent application anti-estrogen drugs could achieve complete tumor control identical to that obtained with continuous MTD treatment. No tumor growth was observed even during a 3-week interval during which therapy was not applied. Advantages of this therapy included a significant (up to 50%) cumulative dose reduction and decreased evidence tumor cell resistance (based on ER and MDR1 expression). Our study does not demonstrate that Tamoxifen intermittent therapy can prolong response as it we have found in chemotherapy for breast cancer in pre-clinical studies and hormone therapy in prostate cancer. Here we were primarily focused on experiments that address a clinical scenario in which the patient presents with highly symptomatic or life-threatening disease requiring rapid and significant reductions of the tumor burden. We demonstrate that such treatment can be administered to substantially reduce the tumor burden while also using interruptions of therapy to reduce evolutionary selection for resistant populations while still maintaining tumor control. Furthermore, these outcomes can be achieved while substantially reducing (by up to 50%) the total dose of Tamoxifen thus reducing toxicity and cost.

Finally, by imaging analysis (radiomics), we have demonstrated that evolutionary-based Tamoxifen therapies develop tumors with the same ER homogeneity than SOC. These results suggest that imaging biomarkers that correlate with intratumoral evolution during treatment may ultimately prove to be useful guides for evolution-based treatments.

## Acknowledgements

This work has been supported in part by the SAIL Core, Tissue Core Facility, the Analytic Microscopy Core Facility, and by the IRAT Core Facility at the H. Lee Moffitt Cancer Center & Research Institute, an NCI designated Comprehensive Cancer Center (P30-CA076292).

---

**Sup. Figure 1:**
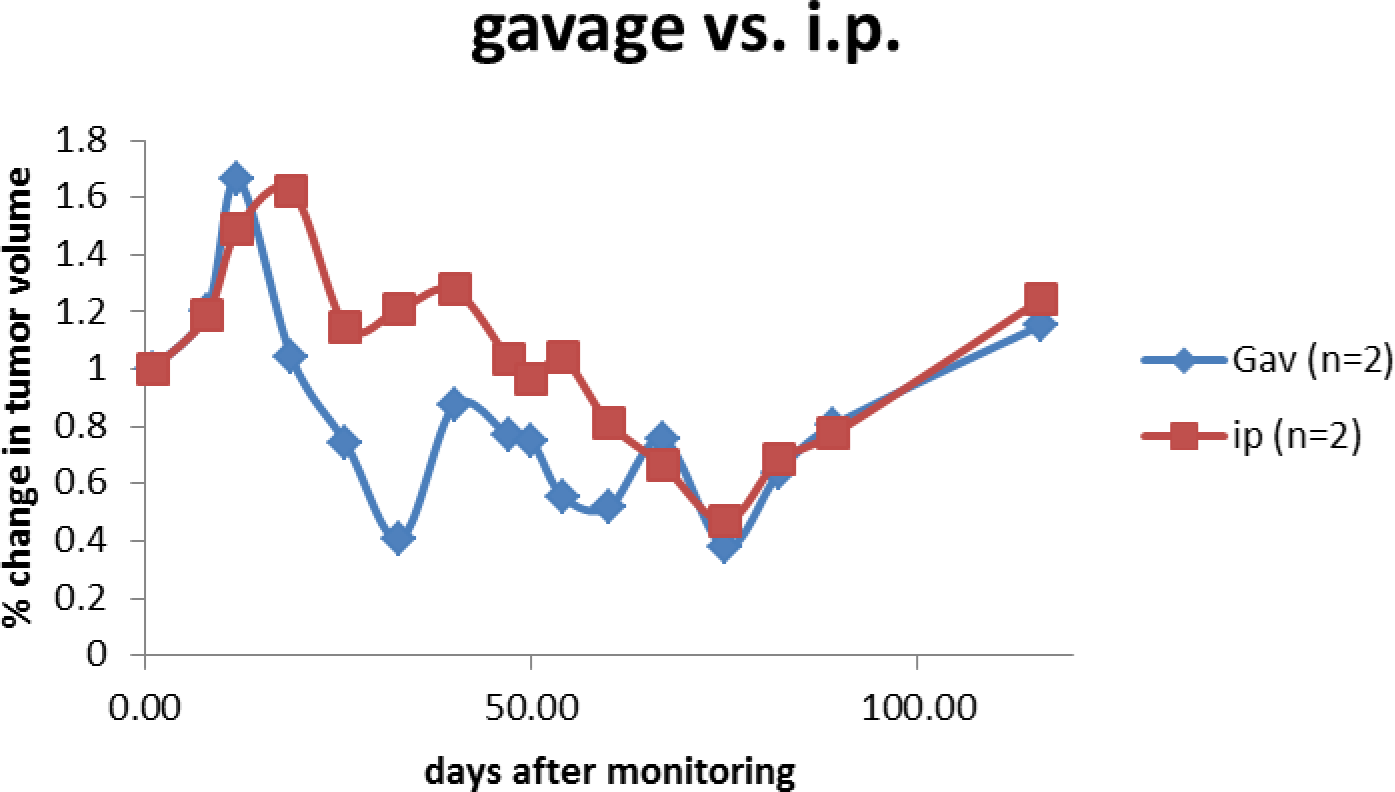
Fold of change in tumor volume in mice treated with tamoxifen by either i.p. injections or gavage.

**Sup. Figure 3:**
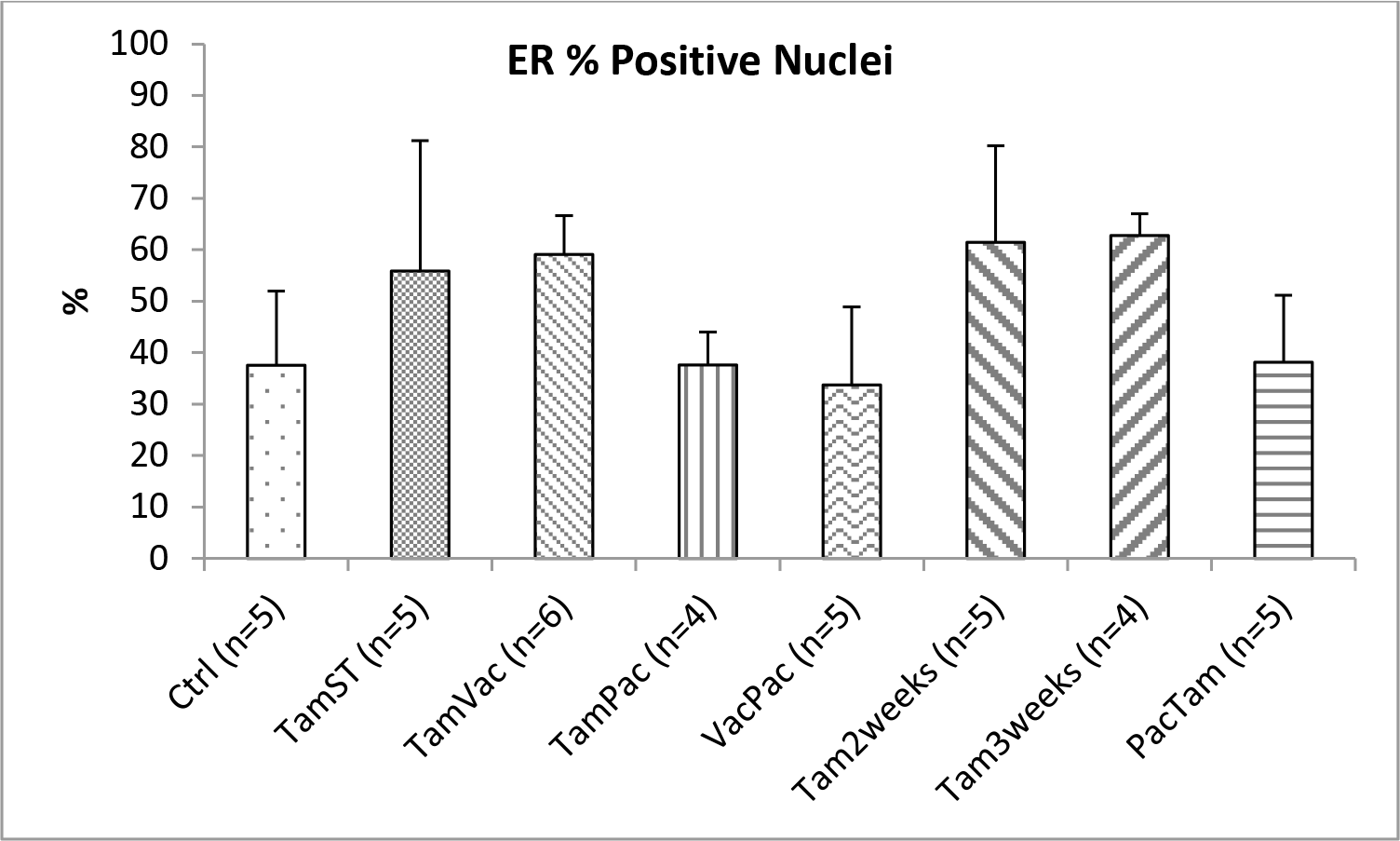
Cohort *C* IHC analysis of the ER expression under different treatments. (Significance calculations were done by using a Student t-test with two-tailed distribution and considering two samples with unequal variance).

**Sup. Figure 4:**
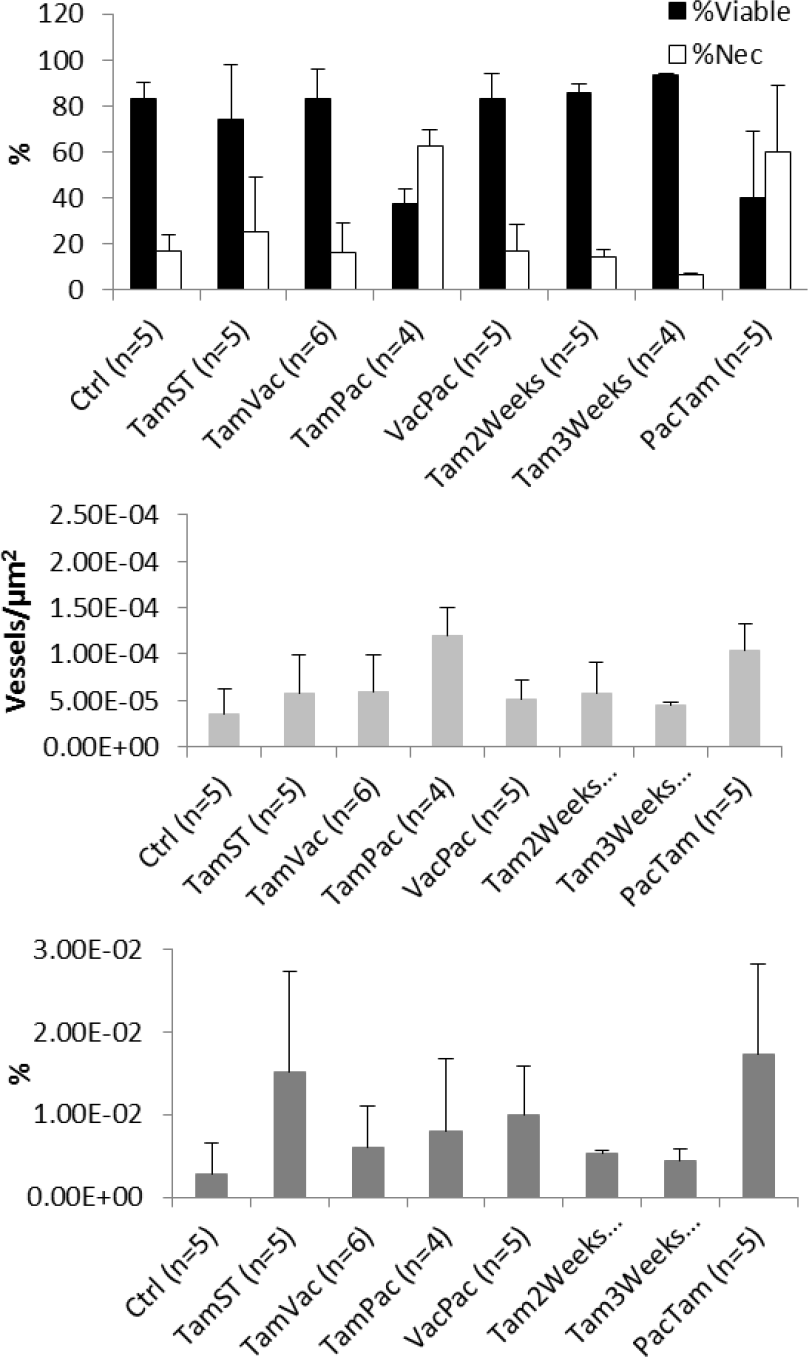
Cohort *C* IHC analysis of viable and necrotic tissues (H&E), vessel density (CD31) and functionality (SMA) in tumors under different treatments.

